# Mul1 suppresses Nlrp3 inflammasome activation through ubiquitination and degradation of Asc

**DOI:** 10.1101/830380

**Authors:** June-Hyung Kim, Yunjong Lee, Gee Young Suh, Yun-Song Lee

## Abstract

Activation of the Nlrp3 inflammasome consisting of three major components, Nlrp3, Asc, and pro-caspase-1, results in the activation of caspase-1 and subsequent proteolytic cleavage of pro-IL-1β and pro-IL-18. To avoid excessive inflammatory response, the Nlrp3 inflammasome has to be precisely controlled. In this study, we show that the mouse mitochondrial E3 ubiquitin protein ligase (Mul1) suppresses Nlrp3 inflammasome activation through ubiquitination and degradation of Asc. In J774A.1 cells, Mul1 overexpression attenuated Nlrp3 activation, whereas Mul1 knockdown augmented Nlrp3 activation in terms of IL-1β secretion and cleavage of pro-caspase-1 and pro-IL-1β. Mul1 interacted with Asc, and ubiquitinated it at K21, K22, K26, and K55 residues, in a K48-linked manner, leading to proteasomal degradation. Convincingly, Mul1-mediated suppression of Nlrp3 activation was inhibited by K21R-, K22R-, K26R-, K52R-Asc mutants in RAW264.7 cells, when compared with the wild-type Asc. Furthermore, Aim2 inflammasome activation was also inhibited by Mul1 in the wild-type Asc-, but not in mutant Asc-expressing RAW264.7 cells. Taken together, these data suggest that Mul1 suppresses Nlrp3 inflammasome activation, through Asc ubiquitination and degradation.

## Introduction

Nlrp3 (NOD-, LRR-, and pyrin domain-containing protein 3) is an intracellular sensor that detects a broad range of microbial pathogens, endogenous danger signals, and environmental irritants [1–4]. Typically, priming by “Signal 1” involves transcriptional and post-transcriptional regulation of related components, such as IL-1β and Nlrp3, through pattern recognition receptors and NFκB signaling. In the following step, recognition of Nlrp3 activator, “Signal 2,” induces the formation of the Nlrp3 inflammasome complex that consists of three major components: Nlrp3, Asc (apoptosis-associated speck-like protein containing a Card; also known as Pycard, Pyd And Card domain containing), and pro-caspase-1, acting as sensor, adaptor, and effector molecules, respectively. Nlrp3, after sensing stimuli, engages multiple molecules of Asc, which, through protein-protein interaction, form oligomers and finally “bird nest”-like supramolecular protein complexes known as Asc specks [5–8]. The supramolecular protein complexes recruit and activate procaspase-1, culminating in the production of active IL-1β and IL-18 through proteolytic cleavage by activated caspase-1.

Since the Nlrp3 inflammasome activation has been implicated in various kinds of diseases, including autoimmune diseases [9, 10], atherosclerosis [11, 12], and type 2 diabetes [13, 14], it needs to be tightly regulated to avoid exaggerated immune responses and a detrimental outcomes [1, 15, 16]. In addition to regulation of NFκB signaling, proteins in the Nlrp3 inflammasome can be controlled by post-translational modification including ubiquitination. For example, Nlrp3 is heavily ubiquitinated under basal conditions and BRCA1/BRCA2-Containing Complex Subunit 3 (BRCC3) promotes de-ubiquitination and activation of Nlrp3 [17], while F-Box And Leucine Rich Repeat Protein 2 (FBXL2) [18] and Tripartite Motif Containing 31 (TRIM31) [19] ubiquitinate Nlrp3 in a K48-linked manner, leading to its proteasomal degradation. Furthermore, cAMP induced through the dopamine receptor D1, binds to Nlrp3 and promotes its K48-linked ubiquitination and phagosomal degradation through Membrane Associated Ring-CH-Type Finger 7 (MARCH7) [20]. IL-1β is also the target for ubiquitination and proteasomal degradation by the endogenous E3 ligase E6AP in human papilloma virus infection [21]. In contrast to Nlrp3 and IL-1β, ubiquitination-mediated negative regulation of Asc remains largely unknown, whereas positive regulation of Asc through linear ubiquitination by Linear Ubiquitin Chain Assembly Complex LUBAC [22] and K63-linked ubiquitination by TNF Receptor Associated Factor 3 (TRAF3) [23] have been reported.

Mitochondrial E3 ubiquitin protein ligase (Mul1 [24], also known as MAPL [25], MULAN [26], or GIDE [27]), is a multifunctional protein that is embedded in the outer mitochondrial membrane with its C-terminal RING finger domain facing the cytoplasm. Mul1 is involved in various physiological processes through its E3 ligase activity, conjugating ubiquitin or small ubiquitin-like modifier (SUMO) to target proteins. For example, Mul1 regulates mitochondrial dynamics through ubiquitination/proteasomal degradation of Mitofusin 2 (Mfn2) [28] and SUMOylation/stabilization of Dynamin Related Protein 1 (DRP1) [29]. Furthermore, Mul1 regulates selenite-induced mitophagy by ubiquitination of Unc-51 Like Autophagy Activating Kinase 1 (ULK1) [30]. In addition to involvement in mitochondrial dynamics and mitophagy, Mul1 controls innate immune response. Mul1 induces SUMOylation and activation of Retinoic Acid-Inducible Gene I (RIG-1), an essential innate immune receptor that binds double-stranded RNA in the cytosol, leading to initiation of antiviral signaling [31]. However, whether and how Mul1 regulates innate immune response in the context of the Nlrp3 inflammasome remains largely unknown.

Here we report a novel regulatory mechanism for the Nlrp3 inflammasome, in which mouse Mul1 suppresses Nlrp3 inflammasome activation through ubiquitination and proteasomal degradation of Asc. Through Mul1 overexpression and knockdown experiments, we demonstrate that Mul1 suppressed Nlrp3 inflammasome activation. Molecularly, Mul1 bound Asc, and mediated K48-linked ubiquitination at multiple sites and subsequent proteasomal degradation. Furthermore, Asc mutants at Mul1-mediated ubiquitination sites inhibited Mul1-mediated suppression of Nlrp3 inflammasome activation. Finally, activation of Aim2 inflammasome, which also uses Asc as an adaptor, was inhibited by Mul1 in J774A.1 cells, while Mul1 did not affect Aim2 inflammasome activation in the mutant Asc-expressing RAW264.7 cells. Taken together, these novel data support the hypothesis that Mul1 may act as a suppressor of the Nlrp3 inflammasome activation through ubiquitination and degradation of Asc.

## Results

### Mul1 negatively regulates Nlrp3 inflammasome activation

First, we addressed whether Mul1 might affect Nlrp3 inflammasome activation in mouse macrophage J774A.1 cells by overexpressing and knocking down Mul1. When J774A.1 cells were infected with the wild-type Mul1-lentivirus, and primed and stimulated with LPS and ATP, respectively, overexpressed Mul1 dramatically suppressed the secretion of IL-1β and cleavage of pro-IL-1β and pro-caspase-1, in a dose-dependent manner, compared with the control lentivirus-infected group (Fig. 1A).

**Figure 1.**
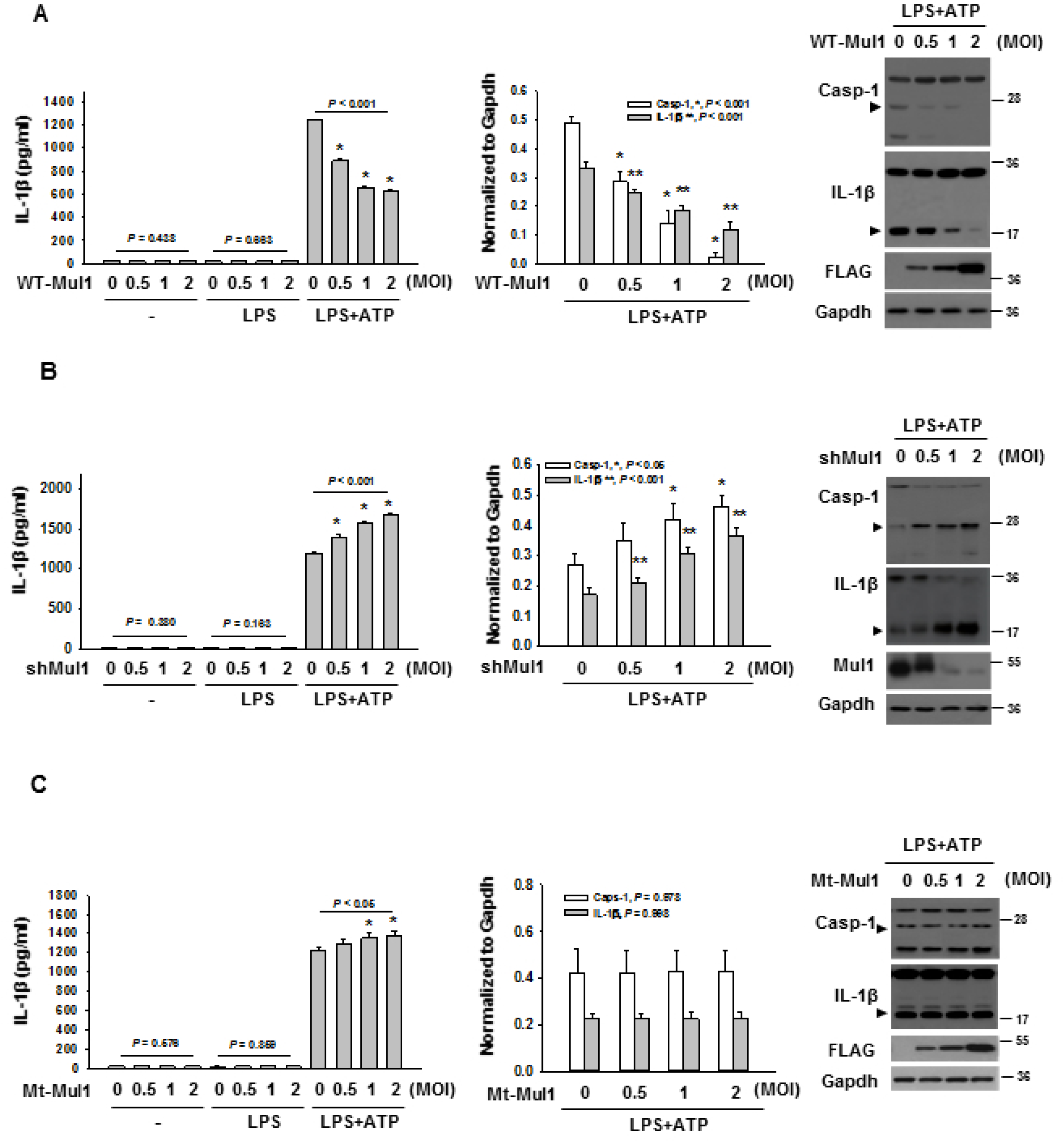
Mul1 negatively regulates the Nlrp3 inflammasome activation induced by LPS and ATP. A A Suppression of Nlrp3 inflammasome activation by Mul1 overexpression. J774A.1 cells were infected with the wild-type Mul1-lentivirus (WT-Mul1) in a 10 cm-dish at 0.5, 1, and 2 MOI (multiplicity of infection). After 2 d, cells were primed with 100 ng/ml LPS for 6 h and stimulated with 5 mM ATP for 1 h. IL-1β secretion into the culture medium and intracellular cleavage of pro-capspase-1 and pro-IL-1β were assessed by ELISA and western blotting, respectively. Arrow heads in representative Western blot images indicate active caspase-1 (p20) and IL-1β (p17). * and ** indicates statistically significant difference, compared with the control lentivirus-infected group (n=4, ANOVA and Holm-Sidak method). B Augmentation of Nlrp3 inflammasome activation by Mul1 knockdown. J774A.1 cells were infected with Mul1 shRNA-lentivirus (shMul1) and assessment of Nlrp3 inflammasome activation was performed as in (A). * and ** indicate statistically significant difference, compared with the control lentivirus-infected group (n=4, IL-1β in ELISA and IL-1β (p17) by ANOVA and Holm-Sidak method and caspase-1 (p20) by ANOVA on ranks and Student-Newman-Keuls method). C Inhibitory effect of Mul1 upon Nlrp3 inflammasome activation depends on the E3 ligase activity of Mul1. Infection of C302/305S-Mul1 (Mt-Mul1)-lentivirus and assessment of Nlrp3 inflammasome activation was performed as in (A). * indicates statistically significant difference, compared with the control lentivirus-infected group (n=4, ANOVA and Holm-Sidak method).

In contrast, Mul1 overexpression did not significantly change TNF-α secretion from J774A.1 cells treated with LPS or LPS+ATP, compared with the control lentivirus-infected group (Appendix Fig. S1A). Moreover, infection of Mul1-lentivirus by itself did not induce secretion of IL-1β and cleavage of pro-IL-1β and pro-caspase-1, in J774A.1 cells treated with LPS alone. Similar to the inhibitory effects on Nlrp3 inflammasome activation induced by LPS and ATP, Mul1 overexpression also significantly suppressed inflammasome activity in LPS-primed J774A.1 cells, stimulated by other Nlrp3 Signal 2 stimuli including nigericin (potassium ionophore), and alum crystal without altering TNF-α secretion (Fig. EV1A and EV2A).

To confirm the inhibitory action of Mul1 on Nlrp3 inflammasome activation, we assessed the effects of Mul1 knockdown using Mul1 shRNA-lentivirus (shMul1). Compared to the control lentivirus, shMu1 significantly augmented IL-1β secretion and cleavage of pro-IL-1β and procaspase-1, in a dose-dependent manner in LPS-primed and ATP-stimulated J774A.1 cells, without any significant changes in TNF-α secretion (Fig. 1B and Appendix Fig. S1B). Similar to shMul1 that was used in the above experiments (derived from TRCN0000040742), another Mul1 shRNA-lentivirus targeting a different sequence of *mul1* (derived from TRCN0000328514) also enhanced IL-1β secretion in J774A.1 cells stimulated with ATP (Appendix Fig. S2), and we used shMul1 derived from TRCN0000040742 for the rest of the experiments. Like the potentiating effects of Mul1 knockdown in Nlrp3 inflammasome activation induced by LPS and ATP, shMul1 also significantly increased secretion of IL-1β and cleavage of pro-IL-1β and procaspase-1, in LPS-primed J774.1 cells stimulated with nigericin and alum, compared with the control lentivirus-infected group (Fig. EV1B and EV2B).

As Mul1 was reported to induce ubiquitination of several proteins, such as p53 [32] and Akt [33], we investigated whether the inhibitory effects of Mul1 on Nlrp3 inflammasome activation might be mediated through the ubiquitin E3 ligase activity by using a RING domain-mutant Mul1 (C302/305S-Mul1, Mt-Mul1, defective in E3 ligase activity) [33]. As shown in Fig. 1C, Mt-Mul1 did not attenuate secretion of IL-1β and cleavage of pro-IL-1β and pro-caspase-1 in LPS-primed- and ATP-stimulated J774A.1 cells. Instead, Mt-Mul1 slightly but significantly increased IL-1β secretion in a dose-dependent manner without any significant change in TNF-α secretion (Fig. 1C and Appendix Fig. S1C). Similarly, Mt-Mul1 caused slight elevation in IL-1β secretion without inhibiting of cleavage of pro-IL-1β and pro-caspase-1 in J774A.1 cells stimulated with nigericin and alum (Fig. EV1C and EV2C, respectively). These results indicated that Mul1 might inhibit Nlrp3 inflammasome activation using the ubiquitin E3 ligase activity of its RING domain, thus prompting us to search for a potential substrate within the Nlrp3 inflammasome components.

### Mul1 reduces Asc levels by K48-linked poly-ubiquitination and subsequent proteasomal degradation

To find a potential substrate for Mul1, we assessed whether Mul1 alters the levels of major Nlrp3 inflammasome components Nlrp3, Asc, and pro-caspase-1 in J774A.1 cells. Interestingly, WT-Mul1 significantly reduced Asc levels in a dose- and time-dependent manner, without noticeable changes in levels of Nlrp3 and pro-caspase-1 (Fig. 2A). In contrast to WT-Mul1, Mt-Mul1 did not reduce Asc levels (Fig. 2B). As Mul1 was reported to be involved in protein ubiquitination/proteasomal degradation as well as mitophagy, we investigated the possible pathway that might contribute to the reduction in Asc levels. When J774A.1 cells overexpressing WT-Mul1 were treated with a proteasomal inhibitor (MG132) and four autophagy inhibitors having different mechanisms (3-methyadenine (3-MA), phosphatidylinositol-3-phosphate kinase inhibitor; choloroquine (CQ), endosomal acidification inhibitor; bafilomycin (Baf), vacuolar H^+^-ATPase inhibitor; leupeptin (Leu), serine/cysteine protease inhibitor), only MG132 notably prevented Asc reduction (Fig. 2C). Furthermore, MG132 inhibited Asc reduction in J774A.1 cells treated with cycloheximide as well as in J774A.1 cells overexpressing Mul1 and treated with cycloheximide. Cycloheximide alone decreased Asc levels and led to a further decrease in Asc in Mul1-overexpressing J774A.1 cells (Fig. 2D).

**Figure 2.**
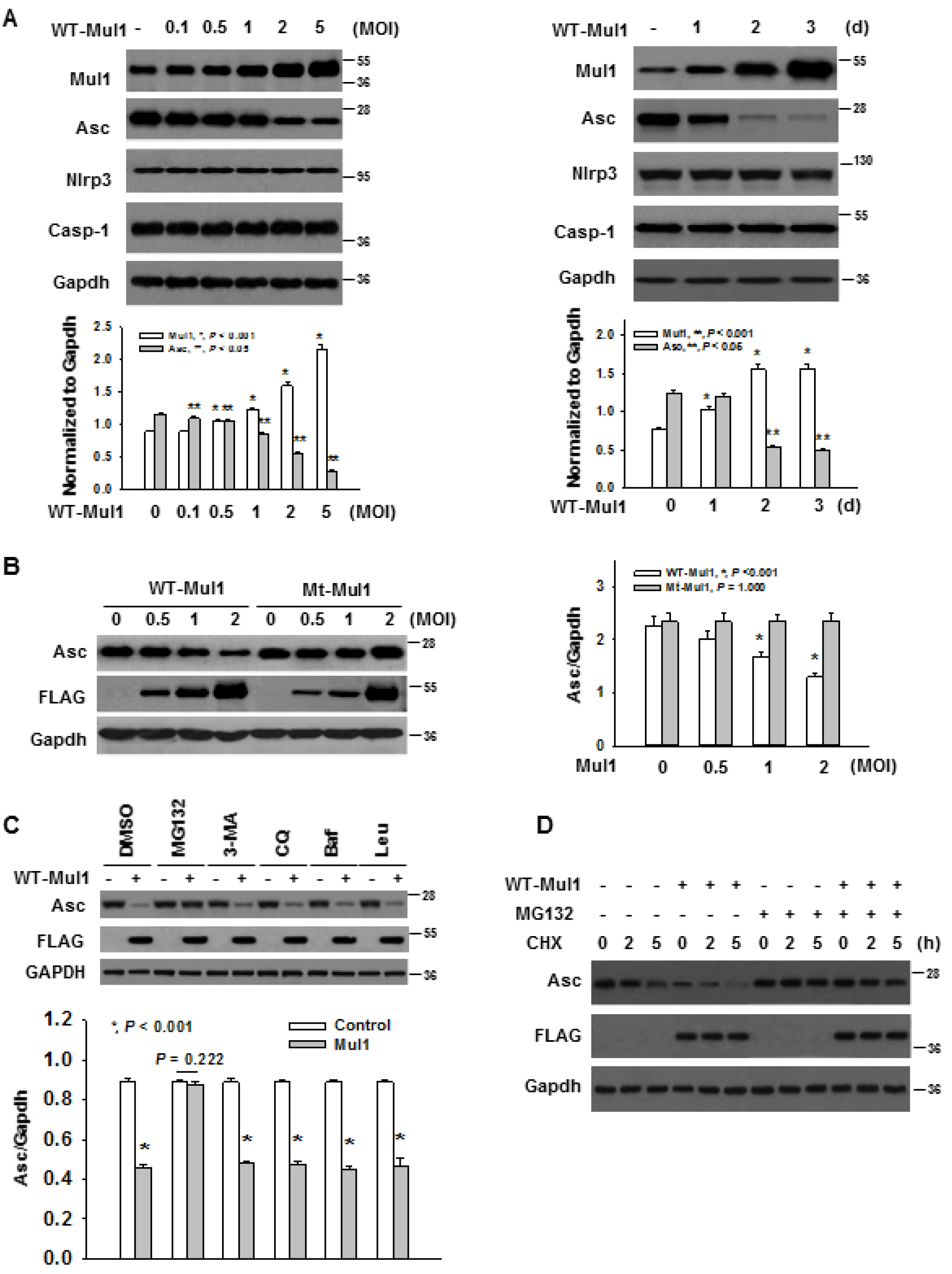
Mul1 induces proteasomal degradation of Asc in J774A.1 cells. A Reduction of Asc levels by Mul1 overexpression in a dose- and time-dependent manner in the left and right panels, respectively. WT-Mul1 was overexpressed as in Figure 1. * and ** indicate statistically significant difference, compared with the control lentivirus-infected group (n=4, Mul1 by ANOVA and Holm-Sidak method and Asc by ANOVA on ranks and Student-Newman-Keuls method). B Dependence of Mul1-mediated Asc reduction on the E3 ligase activity of Mul1. J774A.1 cells were infected with the wild-type (WT-) and C302/305S-(Mt-) Mul1-lentiviruses at doses of 0.5, 1, and 2 MOI. After 2 d, Asc levels in cell lysates were estimated by western blotting. * indicates statistically significant difference, compared with the control lentivirus-infected group (n=4, ANOVA and Holm-Sidak method). C Mul1-mediated Asc reduction through proteasomal degradation. J774A.1 cells were infected with the wild-type Mul1-lentivirus, and treated with inhibitors 2 d after infection; 5 μM MG132 for 16 h; 1 mM 3-methyadenine (3-MA) for 3 h; 10 μM choloroquine (CQ) for 16 h; 0.5 nM bafilomycin (Baf) for 3 h; 50 μM leupeptin (Leu) for 6 h, prior to cell harvest. * indicates statistically significant difference between the absence and presence of Mul1 overexpression (n=4, Student *t*-test). D Cycloheximide (CHX) chase assay. Two days after lentiviral infection, the culture medium was replaced with fresh medium. Thereafter, MG132 was added at 10 μM for 5 h, and then cycloheximide added at 50 μM for indicated periods, prior to cell harvest.

As these data implied Mul1 facilitated Asc degradation via the proteasome, we addressed Mul1-mediated Asc ubiquitination. Using an *in vivo* ubiquitination assay in HEK293 cells, we found that Asc was poly-ubiquitinated by WT-Mul1 in a complete reaction mixture (Fig. 3A), while Mt-Mul1 was unable to do so (Fig. 3B). In the experiment to determine ubiquitin linkage mode, Asc poly-ubiquitination was seen only with the wild-type and K48-ubiquitin (all Lys residues replaced with Arg except for K48) (Fig. 3C). Asc poly-ubiquitination was not seen with K63-ubiquitin (all Lys residues replaced with Arg except for K63), K48R-ubiquitin (K48 replaced with Arg), and KO-ubiquitin (all Lys residues replaced with Arg). Additionally, WT-Mul1 but not Mt-Mul1 recombinant protein induced poly-ubiquitination of Asc *in vitro* (Fig. 3D).

**Figure 3.**
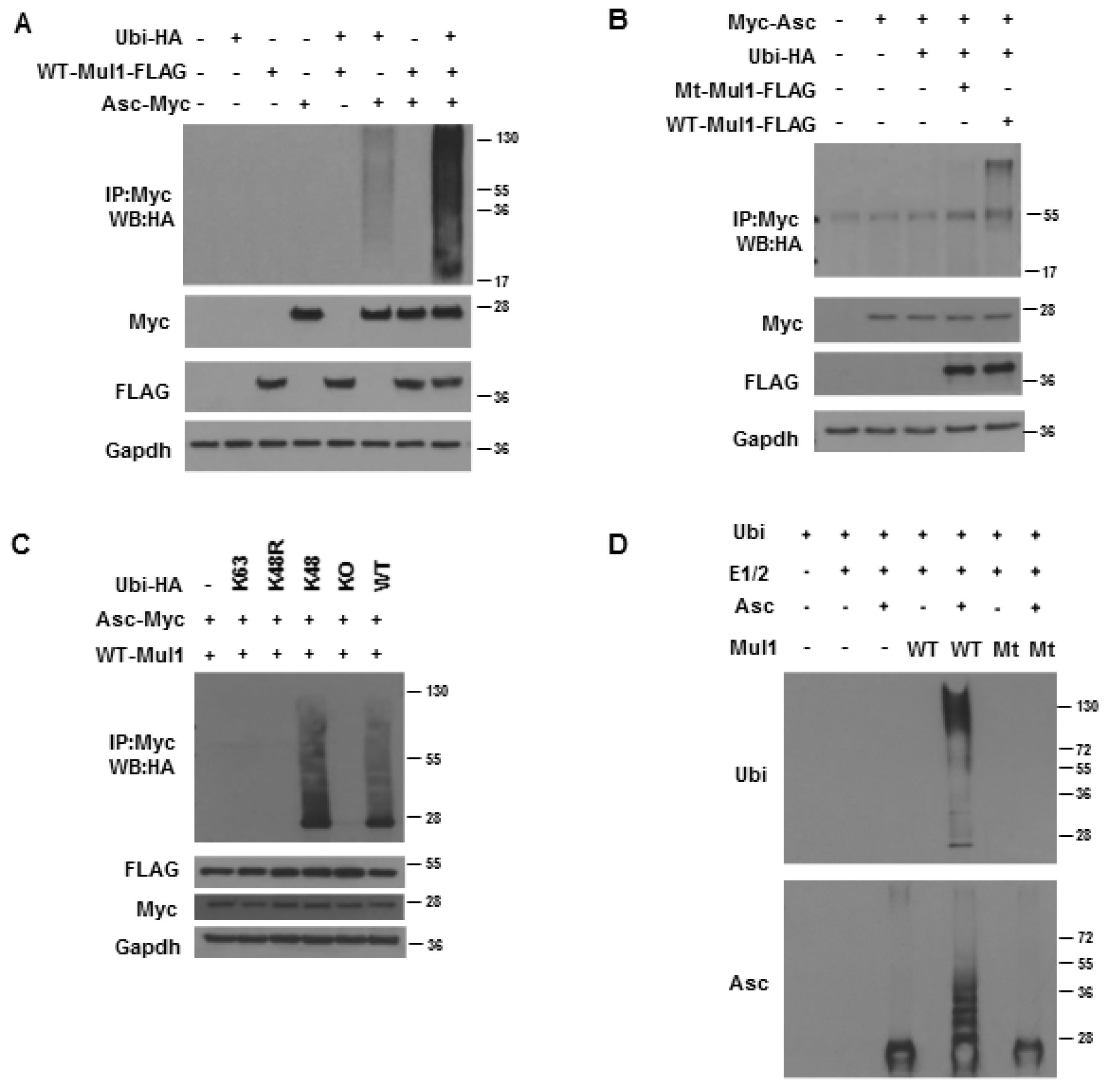
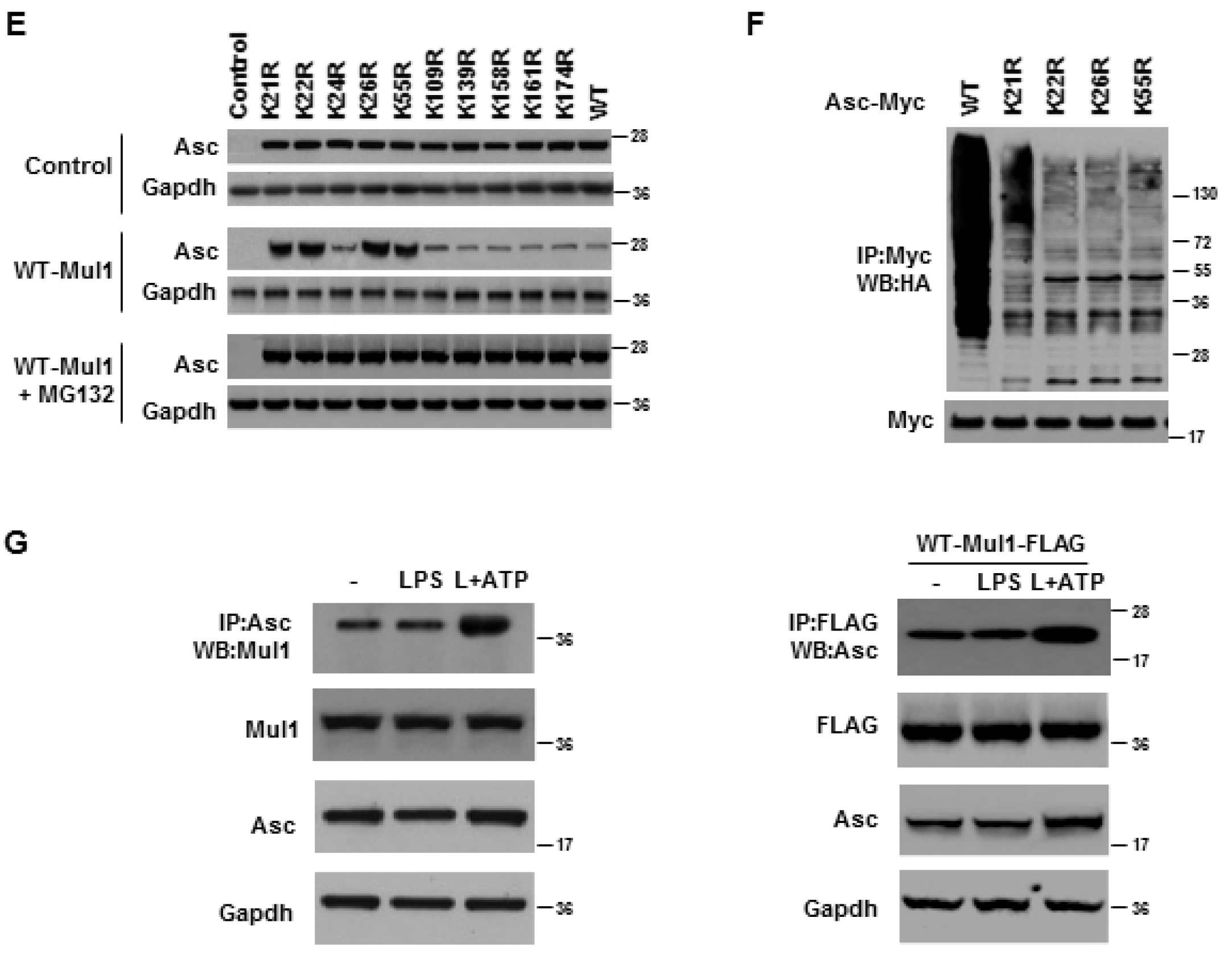
Mul1 induces poly-ubiquitination of Asc. A *In vivo* ubiquitination assay. Mul1-FLAG, Asc-Myc, and hemagglutinin-tagged ubiquitin (Ubi-HA) were expressed in HEK293 cells. Asc was immunoprecipitated (IP) with anti-Myc beads, and ubiquitinated Asc was detected by anti-HA antibody by western blotting (WB). B Requirement of the E3 ligase activity of Mul1 for Asc ubiquitination. *In vivo* ubiquitination assay was performed with the wild-type (WT-) and C302/305S-(Mt-) Mul1 as in (A). C Determination of ubiquitin linkage mode. Ubiquitin linkage mode was determined by using the wild-type and mutant ubiquitin as in (A). WT-Ubi, wild-type ubiquitin; K63-Ubi, all Lys residues replaced with Arg except for K63; K48R-Ubi, K48 replaced with Arg; K48-Ubi, all Lys residues replaced with Arg except for K48; KO-Ubi, all Lys residues replaced with Arg. D *In vitro* ubiquitination assay. Recombinant Asc was incubated with purified wild-type (WT-) or C302/305S-(Mt-) recombinant Mul1, mouse E1/E2, and ubiquitin. Ubiquitinated Asc was detected with anti-ubiquitin and anti-Asc antibodies by western blotting. E Comparison of Mul1-mediated proteasomal degradation of the wild-type and Lys mutant Asc in RAW264.7 cells. Ten K➔R single Asc mutants were expressed in RAW264.7 cells with Mul1, and the cells were harvested after two days. MG132 at 5 μM was added 16 h prior to cell harvest. *Upper*, Asc expression only; *middle*, expression of Asc and Mul1; *lower*, expression of Asc and Mul1, and treatment with MG132. F Comparison of Mul1-mediated ubiquitination of the wild-type and ubiquitination site mutant Asc. *In vivo* Asc ubiquitination assay was performed in HEK293 cells using the wild-type and mutant Asc as in (A). Mul1 and Ubi-HA were included in all lanes. G Interaction between Mul1 and Asc in J774A.1 cells. *Left*, interaction of endogenous Mul1 and endogenous Asc in J774A.1 cells. Mul1 interacting with Asc was detected by co-immunoprecipitation with anti-Asc antibody, followed by western blotting with anti-Mul1 antibody. *Right*; interaction between exogenous Mul1-FLAG and endogenous Asc. Endogenous Asc interacting with Mul1-FLAG was detected by co-immunoprecipitation with anti-FLAG agarose beads, followed by western blotting with anti-Asc antibody.

Next, we determined the site of Mul1-mediated poly-ubiquitination in Asc. For this aim, we generated ten K➔R single mutant Asc plasmids, in which each Lys residue (K21, 22, 24, 26, 55, 109, 139, 158, 161, and 174) was replaced with Arg. When the wild-type (WT-) and mutant Asc plasmids were transfected into RAW264.7 cells that do not to express Asc [34], all Asc mutants were expressed to a similar extent (Fig. 3E, *upper*). In these cells, co-expression of Mul1 reduced the levels of the wild-type and all the Asc mutants, except K21R-, K22R-, K26R-, and K55R-Asc (Fig. 3E, *middle*), and the levels of the wild-type and all the Asc mutants including K21R-, K22R-, K26R-, and K55R-Asc became similar by MG132 treatment (Fig. 3E, *lower*). Consistent with resistance to Mul1-mediated degradation in K21R-, K22R-, K26R-, and K55R-Asc, Mul1-induced poly-ubiquitination was remarkably reduced in K21R-, K22R-, K26R-, and K55R-Asc, compared to WT-Asc in an *in vivo* ubiquitination assay in HEK293 cells (Fig. 3F).

To observe the interaction between Mul1 and Asc, both Mul1-FLAG and Asc-Myc were expressed in HEK293 cells and their interaction was checked by a reciprocal co-immunoprecipitation assay (Fig. EV3). After immunoprecipitation with anti-Myc and anti-FLAG antibodies, Mul1 and Asc were detected by western blotting, respectively. In addition, the interaction of endogenous Mul1 with endogenous Asc in J774A.1 cells was detected by co-immunoprecipitation using an anti-Asc antibody (Fig. 3G, *left*). Furthermore, the binding of endogenous Asc to Mul1 was also observed in J774.1 cells expressing Mul1-FLAG (Fig. 3G, *right*). Interestingly, the interaction between Mul1 and Asc was also observed under basal and LPS-primed conditions, and their interaction was enhanced by ATP stimulation, compared with the basal and LPS primed conditions.

### Mutation of Mul1-mediated ubiquitination sites in Asc attenuates Mul1-mediated suppression of IL-1β secretion from LPS-primed and ATP-stimulated RAW264.7 cells

Next, we sought to confirm whether Mul1-mediated suppression of Nlrp3 inflammasome activation might be mediated by ubiquitination and proteasomal degradation of Asc. For this aim, IL-1β secretion was measured in RAW264.7 cells expressing WT-, K21R-, K22R-, K26R-, and K55R-Asc. As seen in Fig. 4A, LPS and ATP clearly induced IL-1β secretion in RAW264.7 cells transfected the wild-type Asc plasmid and was not observed in cells transfected with the control plasmid, as expected. Compared with WT-Asc, K21R-, K22R-, K26R-, and K55R-Asc slightly but significantly elevated IL-1β secretion upon ATP stimulation. Moreover, Mul1 overexpression did not suppress IL-1β secretion from RAW264.7 cells expressing K21R-, K22R-, K26R-, and K55R-Asc, while on the other hand, Mul1 overexpression remarkably attenuated IL-1β secretion from RAW264.7 cells expressing WT-Asc. The interference of Mul1-mediated suppression of IL-1β secretion by mutation of Mul1-mediated ubiquitination sites in Asc does not seem to be derived from the reduced binding of Mul1 to the Asc mutants, because similar binding of WT-, K21R-, K22R-, K26R-, and K55R-Asc to Mul1 was observed in a co-immunoprecipitation assay (Fig. 4B). Using amino acid sequence alignment analysis, we noticed that K21, K22, K26, and K55 of the pyrin domain (Pyd) in mouse Asc were conserved in human, chimpanzee (*Pan troglodytes)*, and rat proteins (Fig. 4C).

**Figure 4.**
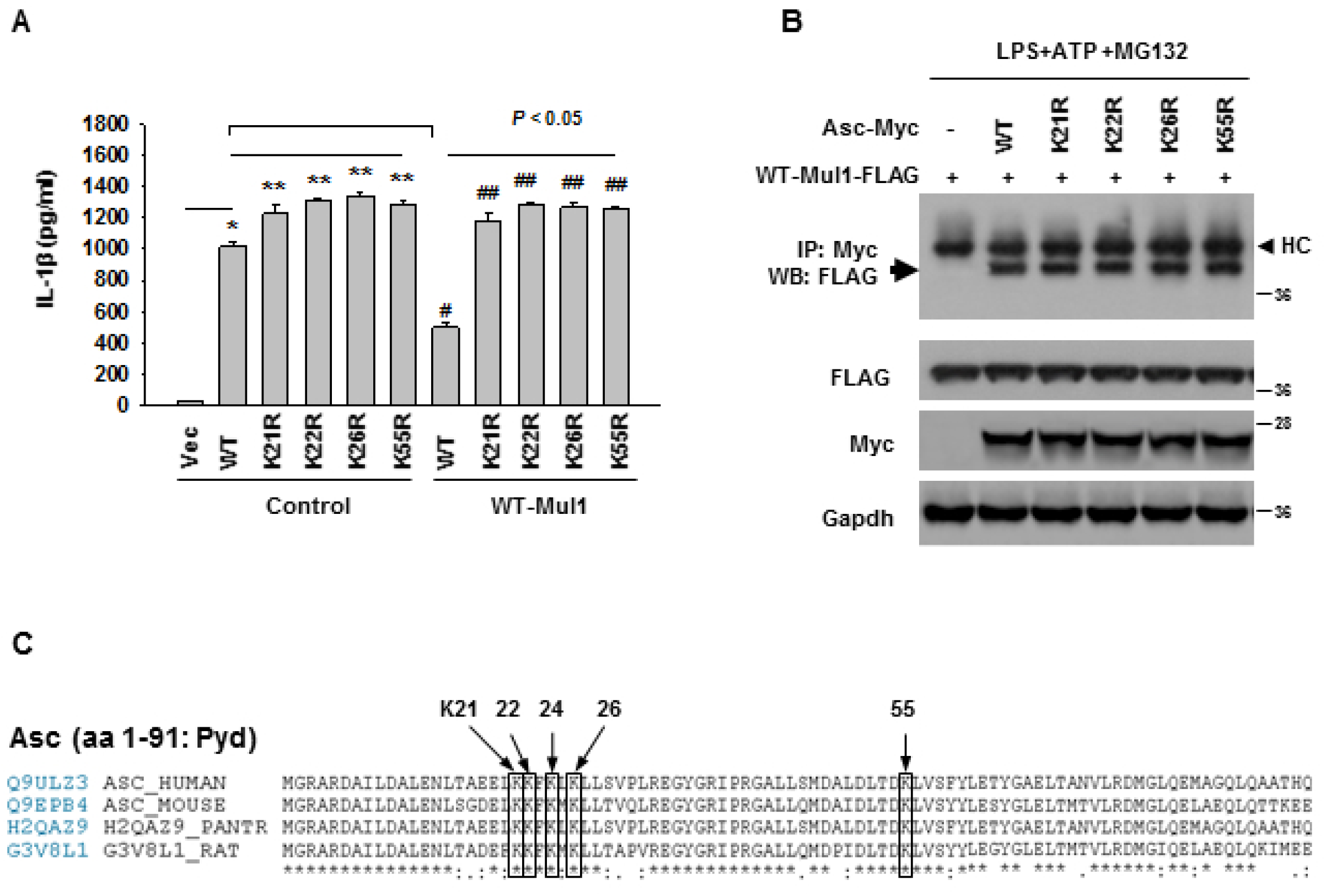
Mutation of Mul1-mediated ubiquitination sites in Asc attenuates Mul1-mediated suppression of IL-1β secretion. A Comparison of IL-1β secretion in RAW264.7 cells expressing the wild-type and mutant Asc. RAW264.7 cells expressing the wild-type or ubiquitination site mutant Asc with or without overexpression of WT-Mul1 were primed and stimulated with LPS and ATP, and IL-1β secretion was assessed by ELISA. Symbols indicate statistically significant difference (n=4 by ANOVA on ranks and Student-Newman-Keuls method). *, compared with control lentivirus; ** and ^#^, compared with WT-Asc only; ^##^, compared with WT-Asc+Mul1. B Comparison of interaction of Mul1 with the wild-type and ubiquitination site mutant Asc. RAW264.7 cells were transfected with the wild-type or mutant Asc together with Mul1, and then primed and stimulated with LPS and ATP. MG132 at 5 μM was added in during LPS priming. Interaction between Mul1 and Asc was detected by co-immunoprecipitation assay as in Figure 3G. HC, heavy chain of IgG. C Conserved Mul1-mediated ubiquitination sites in the pyrin domain (Pyd) of mammalian Asc. Alignment analysis of the amino acid sequence was performed by the comparison tool in UniProt (www.uniprot.org).

### Mul1 suppresses Aim2 inflammasome activation

As Asc also acts as an adaptor for other types of inflammasome complexes including the Aim2 inflammasome [35], we addressed whether Mul1 might play a suppressive role in Aim2 inflammasome activation by Asc ubiquitination. Similar to the suppressive effects of Mul1 upon Nlrp3 inflammasome activation seen in Fig. 1, Mul1 overexpression significantly attenuated secretion of IL-1β and cleavage of pro-caspase-1 and pro-IL-1β in a dose-dependent manner in J774A.1 cells primed with LPS and stimulated with poly(dA:dT) (Fig. 5A), while Mul1 knockdown augmented secretion of IL-1β and cleavage of pro-caspase-1 and pro-IL-1β (Fig. 5B). In addition, Mt-Mul1 did not suppress the secretion of IL-1β and cleavage of pro-caspase-1 and pro-IL-1β (Fig. 5C). Instead, Mt-Mul1 slightly but significantly augmented IL-1β secretion upon poly(dA:dT) stimulation. Furthermore, K21R-, K22R-, K26R-, and K55R-Asc annulled Mul1-mediated suppression of IL-1β secretion and cleavage of pro-caspase-1 and pro-IL-1β in RAW264.7 cells stimulated with poly(dA:dT) (Fig. 5D).

**Figure 5.**
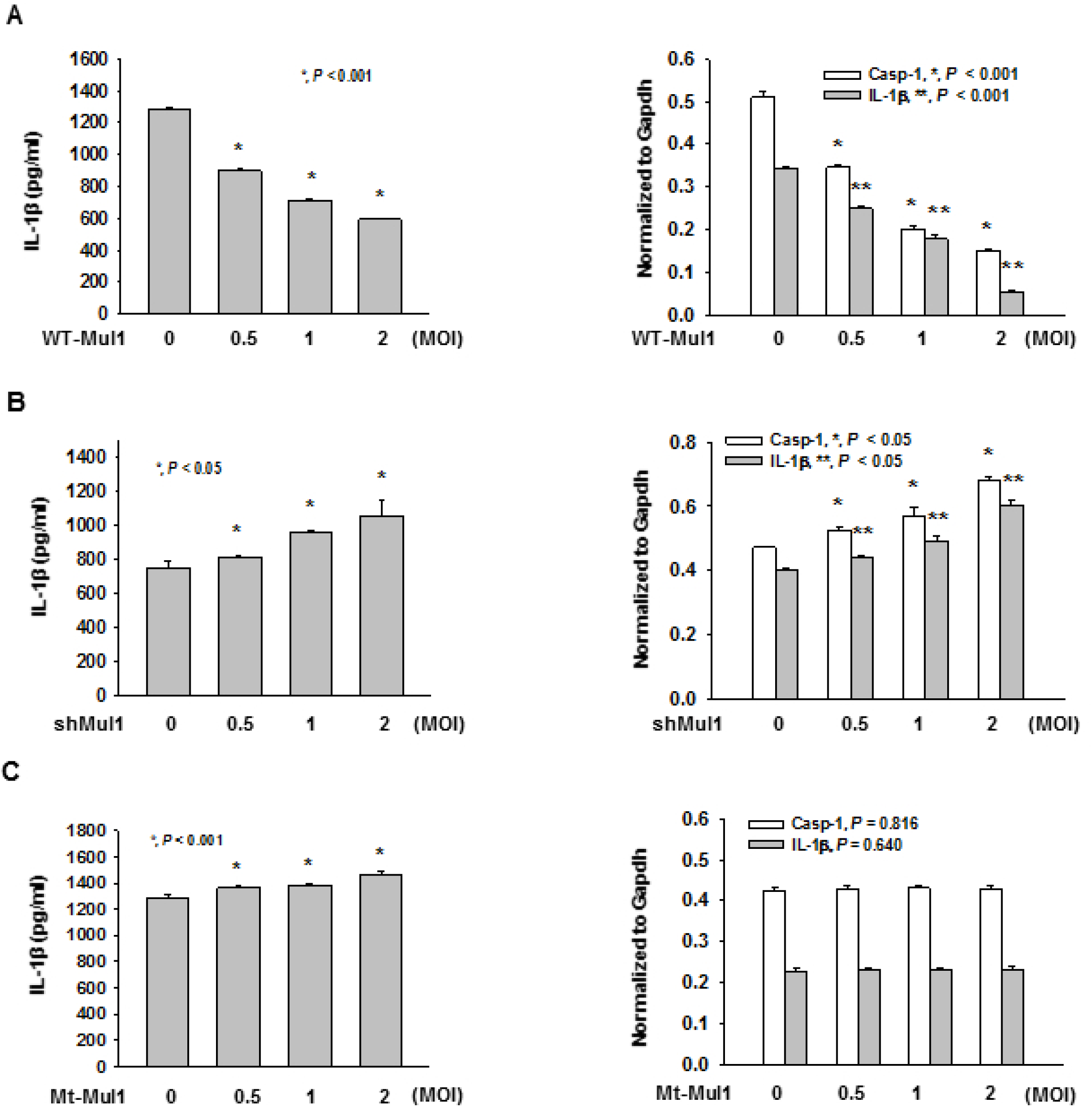
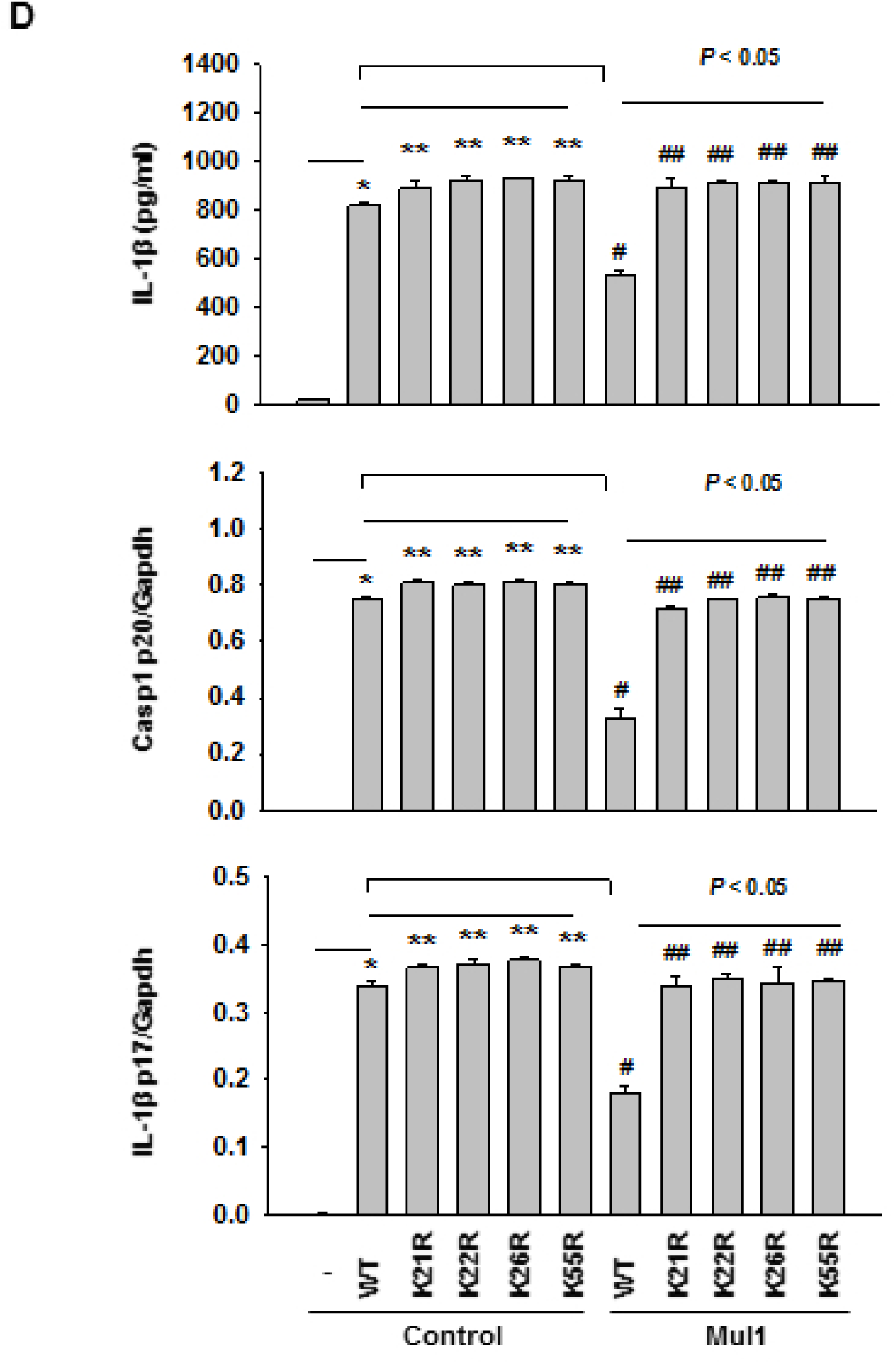
Mul1 suppresses the Aim2 inflammasome activation. Experiments performed as in Figure 1, except Aim2 inflammasome activation by LPS and poly(dA:dT). Aim2 inflammasome activation was induced by transfecting LPS-primed J774A.1 cells with poly(dA:dT) 6 h prior to cell harvest. A Suppression of Aim2 inflammasome activation by Mul1 overexpression. * and ** indicate statistically significant difference, compared with the control lentivirus-infected group (n=4, ANOVA and Holm-Sidak method). B Augmentation of Aim2 inflammasome activation by Mul1 knockdown. * and ** indicate statistically significant difference, compared with the control lentivirus-infected group (n=4, ANOVA on ranks and Student-Newman-Keuls method). C Inhibitory effect of Mul1 on Aim2 inflammasome activation is dependent on the E3 ligase activity of Mul1. Infection of C302/305S-Mul1 (Mt-Mul1)-lentivirus and assessment of Aim2 inflammasome activation was performed as in (A). * indicates statistically significant difference, compared with the control lentivirus-infected group (n=4, ANOVA and Holm-Sidak method). D Attenuation of Mul1-mediated suppression of Aim2 inflammasome activation by mutation of Mul1-mediated ubiquitination sites in Asc in RAW264.7 cells. Symbols indicates statistically significant difference (n=4 by ANOVA on ranks and Student-Newman-Keuls method). *, compared with control lentivirus; ** and ^#^, compared with WT-Asc only; ^##^, compared with WT-Asc+Mul1.

## Discussion

In this study, we clearly demonstrated that Mul1 suppresses the Nlrp3 inflammasome by promoting ubiquitination and degradation of Asc, an adaptor of the Nlrp3 inflammasome complex. We supported this hypothesis by showing 1) suppression and augmentation of Nlrp3 inflammasome activation by Mul1 overexpression and knockdown, respectively, 2) direct Asc ubiquitination by Mul1 *in vivo* and *in vitro*, 3) resistance of ubiquitination and degradation of the Asc mutants to Mul1, and 4) interference of Mul1-mediated suppression of Nlrp3 inflammasome activation by Asc mutants. Furthermore, similar observations with the Aim2 inflammasome activation supports the idea that Asc may be a target for Mul1 in suppression of inflammasome activation.

Asc is composed of an N-terminal Pyd, an unstructured linker region, and a C-terminal Card [1-4, 6, 7]. Once inflammasome sensors, that typically contain a Pyd, are activated by recognizing inflammasome activators, they act as a seed for the recruitment of Asc through Pyd:Pyd homotypic interactions between inflammasome sensors and Asc, leading to the formation of Asc oligomers. Asc oligomers, in turn, nucleate filamentous polymers of Asc through inter-strand Card:Card interactions in Asc, forming Asc specks that are micrometer-sized supramolecular protein complexes resembling a bird’s nest. Asc specks are the sites of pro-caspase-1 recruitment and activation through Card:Card interactions between Asc and pro-caspase-1, and proteolytically activate pro-IL-1β, pro-IL-18, and pore-forming gasdermin D that induces pyroptosis, a type of pro-inflammatory cell death. In addition to the intracellular role of Asc specks, these complexes are released into the extracellular space, and amplify the inflammatory response by activating extracellular pro-caspase-1 and pro-IL-1β and acting as a seed for new Asc speck formation after being engulfed by bystander macrophages [36, 37].

In this context, Asc plays a key role in inflammasome activation, and therefore may be a critical target for precise and appropriate control of inflammasome activation. Post-transcriptional modifications including phosphorylation and ubiquitination have been reported to regulate Asc. Asc is phosphorylated during Nlrp3 and Aim2 inflammasome activation by JNK and Syk, and Syk-mediated phosphorylation at Tyr144 residue in the Card domain of mouse Asc is critical for Asc speck formation and caspase-1 activation [38]. IKKα negatively regulates Asc by restraining Asc in the nuclei, and Signal 2 of Nlrp3 activation recruits protein phosphatase 2 and inhibits IKKα, thus allowing Asc to participate in the Nlrp3 inflammasome activation [39]. In addition to phosphorylation, ubiquitination of Asc is a known regulatory mechanism. LUBAC, consisting of HOIL-1L, HOIP, and SHARPIN, induces linear ubiquitination of Asc that enables Nlrp3 inflammasome activation [22]. Additionally, mitochondrial antiviral signaling protein (MAVS) was reported to stabilize Asc by recruiting TRAF3 that promoted K63-linked ubiquitination of Asc, and enhance Asc specks and IL-1β secretion upon infection with the RNA virus vesicular stomatitis virus [23]. However, the MAVS-TRAF3-Asc axis did not work with other Nlrp3 activators including monosodium urate and calcium pyrophosphate dehydrate. In contrast, Shi et al. reported that Asc aggregates are poly-ubiquitinated in a K63-linked manner and suggested that poly-ubiquitinated inflammasomes can be trafficked by p62 into the autophagic pathway, thus limiting Aim2 inflammasome activation [40]. However, they did not characterize the ubiquitin E3 ligase involved, the ubiquitination sites in Asc, and the causal relationship between Asc ubiquitination and inflammasome activation. To the best of our knowledge, K48-linked ubiquitination and proteasomal degradation-mediated negative regulation of Asc has not been reported yet.

In this study, we demonstrated that Mul1 negatively regulated both Nlrp3 and Aim2 inflammasome activation. We envisage that additional kinds of inflammasomes including NLRC4 and pyrin inflammasomes may be regulated by Mul1, because they use Asc as an adaptor for the inflammasome complex formation as well [41, 42]. In the case of NLRP1b inflammasome activation, while Asc is dispensable for IL-1β secretion and pyroptosis, it is required for Asc speck formation and subsequent activation of caspase-1, which augments IL-1β cleavage [43]. Hence, NLRP1b inflammasome activation may be partially inhibited by Mul1, if Mul1 is engaged in the negative control of NLRP1b inflammasome activation through Asc ubiquitination and degradation. Another emergent question is whether Mul1 suppresses inflammasome activation through ubiquitination and degradation of single Asc molecules or Asc in oligomers and specks. As the ubiquitination sites (K21, K22, K26, and K55) in Asc reside in the Pyd, the Pyd:Pyd homotypic interaction in Asc oligomers may sterically hinder binding of Mul1. Furthermore, the chance for Mul1 to access to Asc may be reduced once Asc specks have formed, because of the compactness of these supramolecular protein complexes. However, we observed increased binding of Mul1 to Asc upon stimulation by Signal 2, and it remains to be elucidated whether Mul1 can bind to Asc oligomers before Asc speck formation or bind single Asc molecules that are translocated into the vicinity of the mitochondria [44]. Furthermore, it needs to be investigated whether other types of post-translational modification of Asc including phosphorylation, linear ubiquitination, and K63-linked ubiquitination may affect Mul1 function by interfering with the Mul1 and Asc interaction.

Mul1 is a multifunctional protein ligase that conjugates ubiquitin or SUMO to substrate proteins, playing distinct biological roles. Mul1 retards cell growth by K48-linked poly-ubiquitination and degradation of Akt [33], while it attenuates apoptosis by K48-linked poly-ubiquitination and degradation of exonuclear p53 and subsequent suppression of p53-dependent transcription [32]. Also, Mfn, which causes mitochondrial hyperfusion, was reported to be a substrate for Mul1 by two research groups. Even though both groups reported that Mfn is poly-ubiquitinated by Mul1, they proposed different scenarios of Mul1 action related to the PINK1/PARKIN pathway for mitophagy. Yun et al. suggested the parallel action of Mul1 and PINK1/PARKIN on Mfn for controlling mitochondrial dynamics [45]. In contrast, Puri et al. proposed that Mul1 may act as a checkpoint for the recovery of mildly stressed mitochondria by degradation of Mfn2, thereby restraining PARKIN from inducing mitophagy [46]. Under severe stress, Mfn2 may impair endoplasmic reticulum (ER)-mitochondria contact, which increases intracellular Ca^2+^ levels, activates Drp1 and mitochondrial fragmentation, and finally induces mitophagy by the PARKIN pathway. In addition to ubiquitination, Mul1 was reported to be involved in mitochondrial dynamics and immune response through its SUMO E3 ligase activity. Mul1 stimulates mitochondrial fission by SUMOylation of Drp1, which then stabilizes ER-mitochondria contact sites that are important for mitochondrial constriction, calcium flux, and cytochrome C release, leading to apoptosis [29]. Doiron et al. found that Mul1 is responsible for antiviral signaling in Sendai virus infection by SUMOylating and subsequently activating RIG-1 [47]. However, this data is contradictory to the previous report by Jenkins et al. that suggested that Mul1 negatively regulated the antiviral response to Sendai virus [31]. Even though the authors proposed Mul1-mediated SUMOylation of RIG-1 as a mechanism for the inhibitory role of Mul1, further study seems to be needed to support that claim.

Recently, Barry et al. clearly demonstrated that Nlrp3 is SUMOylated by Mul1 and Signal 2-dependent de-SUMOylation of Nlrp3 by the SUMO-specific proteases Senp2 and Senp7 promote Nlrp3 activation [48]. In contrast to the inhibitory action of Mul1 on Nlrp3 inflammasome activation, the authors showed that Mul1 did not suppress Aim2-mediated caspase-1 activation in response to cytosolic DNA, even though they observed that Mul1 depletion enhanced Asc oligomerization. This data is contradictory to our findings that Mul1 overexpression suppressed Aim2 inflammasome activation, while Mul1 knockdown had the opposite effect. Moreover, the data obtained from our experiments using the Asc mutants in RAW264.7 cells support the idea that Mul1 can inhibit Aim2 inflammasome activation through Asc ubiquitination, similar to that seen with the Nlrp3 inflammasome activation. The discrepancy of the role of Mul1 in Aim2 inflammasome activation may have originated from different experimental conditions. Barry et al. used primary bone marrow-derived macrophages (BMDMs), and we used the J774A.1 cell line. Another reason for the discrepancy might be the efficiency of Mul1 knockdown. Barry et al., reported that Mul1 knockdown efficiency was around 50% in BMDMs, which is lower than the efficiency in our experiments (Fig. 1 & 5, and Appendix Fig. S3). Moreover, because they did not check the effects of Mul1 overexpression on the Aim2 inflammasome activation, their claim that Mul1 may not be involved in the regulation of Aim2, seems to be further examined. Even with this discrepancy, it is of note that Mul1-mediated Nlrp3 SUMOylation disappeared upon stimulation by Signal 2 in the above report, and Mul1-Asc binding was noticeably increased by Signal 2 in our experiments. Hence, the scenario for dual roles of Mul1 may be plausible that Mul1 may switch its role from Nlrp3 SUMOylation under basal conditions to Asc ubiquitination/degradation under Nlrp3 stimulated conditions.

Even with the difference in the biochemical roles of Mul1, both roles of Mul1 may be physiologically beneficial in the context of regulation of Nlrp3 inflammasome activation, and Mul1 induction can be a therapeutic strategy for inhibiting excessive Nlrp3 inflammasome activation. Till now, transcriptional regulation of Mul1 remains largely unknown, and only forkhead box O (FOXO) transcriptional factors were reported to be involved in Mul1 expression. Cisplatin induced Mul1 via FOXO3, leading to Akt ubiquitination and cell death [49], and Mul1 expression was inhibited by transfection of peroxisome proliferator-activated receptor γ-coactivator 1α (PGC-1α), which not only suppressed FOXO1 and FOXO3, but also attenuated Mfn2 ubiquitination and degradation [50]. As FOXO3 is directly and indirectly activated by AMP-activated protein kinase (AMPK), a master regulator of metabolism and mitochondrial homeostasis [51], the AMPK/FOXO pathway may be a good candidate for Mul1 induction. In support of this idea, the anti-diabetic drug metformin that activates AMPK [52] and reduced Nlrp3 inflammasome activation [13] induced Mul1 expression [53].

Damaged mitochondria play a key role in inflammasome activation not only by generating harmful Nlrp3 activators including oxidized mitochondrial DNA [54] and reactive oxygen species [44], but also by acting as a platform for the Nlrp3 inflammasome assembly through binding of Nlrp3, Asc, and pro-caspase-1 to mitochondrial cardiolipin, which is externalized to the outer mitochondrial membrane at priming [55, 56]. Hence, mitophagy is considered to be an important process for limiting the Nlrp3 inflammasome activation. Drp1 and Mfn control mitochondrial dynamics by inducing mitochondrial fission and fusion, respectively, are suggested to play a role in the Nlrp3 inflammasome activation. RNA virus infection promotes activation of Drp1 by the RIP1-RIP3-dependent cascade resulting in mitochondrial damage and Nlrp3 inflammasome activation [57], whereas Park et al. reported that Drp1 knockdown led to increased Nlrp3 inflammasome activation by ATP and nigericin [58]. As Drp1 is a substrate for Mul1-mediated SUMOylation, it needs to be assessed whether Mul1 may regulate inflammasome activation through Drp1 SUMOylation. In addition to SUMOylation, Ichinohe et al., found that Mfn2 is required for Nlrp3 inflammasome activation, proposing a model wherein Nlrp3 and MAVS associate with Mfn2 and recruit Asc and pro-caspase-1 to the outer mitochondrial membrane [59]. Mul1 promotes Mfn2 ubiquitination and proteasomal degradation, leading to mitochondrial fragmentation and mitophagy. Thus, we cannot exclude the possibility that Mul1-mediated Mfn2 reduction may contribute to suppression of Nlrp3 inflammasome activation, even though we have demonstrated that Asc ubiquitinated by Mul1 was destined for proteasomal degradation, independently of autophagy (Fig. 2).

In summary, we demonstrate that Mul1 suppresses Nlrp3 inflammasome activation through ubiquitination and degradation of Asc in macrophages. We think that additional experiments are needed to elucidate several questions. Does Mul1 deficiency or overexpression affect the inflammasome activity in animal models of inflammasome activation? Is mitophagy an additional mechanism for Mul1-mediated suppression of inflammasome activation? Is Mul1-mediated Asc ubiquitination and degradation affected by other Asc modification including phosphporylation and linear or K63-linked ubiquitination? Does Mul1 target other proteins involved in inflammasome activation? Finally, it is needed to look for an Mul1 inducer and understand its molecular mechanism to development a therapeutic strategy for controlling inflammatory responses.

## Materials and Methods

### Materials

Lipopolysaccharide (LPS from *Escherichia coli*, 055:B5), ATP, cycloheximide, MG132, 3-methyladenine, chloroquine, bafilomycin, and leupeptin were purchased from Sigma (St. Louis, MO, USA). Nigericin and poly(dA:dT)/LyoVec were from Invivogen (San Diego, CA, USA), and alum from Thermo Fisher (Waltham, MA, USA). Antibodies were obtained from the following sources: anti-Mul1 and anti-caspase-1 (Abcam, Cambridge, MA, USA), anti-IL-1β, anti-Nlrp3, anti-Asc, horse radish peroxidase (HRP)-conjugated anti-rabbit IgG, and anti-mouse IgG (Cell Signaling, Beverly, MA, USA), anti-Asc (Proteintech, Rosemont, IL, USA), anti-ubiquitin (Santa Crus, Dallas, TX, USA), anti-hemagglutinin (HA), anti-FLAG, and anti-Myc (Sigma), and anti-glyceraldehyde 3-phosphate dehydrogenase (Gapdh, AbFrontier, Seoul, Korea). Anti-FLAG (M2) and anti-Myc agarose beads were purchased from Sigma and Thermo Fisher (Waltham, MA, USA), respectively. Detailed information about antibodies can be found in Appendix Table 1.

### Cell culture and activation of Nlrp3 and Aim2 inflammasomes

J774A.1, RAW264.7, and HEK293 cells were from the Korea Cell Line Bank (Seoul, Korea), and HEK293T cells from the American Type Culture Collection (Manassas, VA, USA). J774A.1 and RAW264.7 cells were grown in RPMI1640 (Sigma), and HEK293 and HEK293T cells in DMEM (Sigma) containing 10% fetal bovine serum (FBS) (Gibco, Gaithersburg, MD, USA). For Nlrp3 inflammasome activation, J774A.1 and RAW264.7 cells in a 10 cm-dish were primed with 100 ng/ml LPS for 6 h and stimulated with 5 mM ATP, 10 μM nigericin, and 100 μg/ml alum for 1 h [60, 61]. Aim2 inflammasome activation in LPS-primed J774A.1 cells was induced by transfection of 5 μg of poly(dA:dT) using Lipofectamine 2000 (Thermo Fisher) 6 h prior to sample collection [35]. After inflammasome activation, culture media and cells were harvested for ELISA and western blot, respectively. Levels of IL-1β and TNF-α in 200 μl culture medium were measured by ELISA kits according to manufacturer’s instructions (DY401 for IL-1 β and DY410 for TNF-α, R&D systems, Minneapolis, MN, USA). Cleavage of pro-IL-1β and pro-caspase-1 was assessed by western blot of 50 μg of protein extracts of cell lysates, based on our previous report [62]. Images of cleaved caspase-1 (p20) and IL-1β (p17) in the western blot were quantified using a software (ImageJ, https://imagej.net).

### Plasmids and lentiviruses

Plasmids of the wild-type mouse Mul1 (pMul1-FLAG) and Asc (pAsc-Myc) were constructed using pCMV6 Mul1-Myc/DDK (MR205346, Origene, Rockville, MD, USA) and pcDNA3-N-FLAG-Asc (a gift from Bruce Beutler, #75134, Addgene, Watertown, MA, USA) as templates, respectively. Plasmids of RING domain mutant Mul1 (C302/305S-Mul1, MT-Mul1) [33] and K➔R mutant Asc were generated by PCR-based mutagenesis using pMul1-FLAG and pAsc-Myc, respectively. To produce lentiviruses, pMul1-FLAG and pAsc-Myc were subcloned into pLenti X2 Hygro DEST (a gift from Eric Campeau and Paul Kaufman, w17-1, #17295, Addgene) and transfected into HEK293T cells together with the helper plasmid (psPAX2, #12260, a gift from Didier Trono, Addgene) and the envelope plasmid (pMD2.G, #12259, a gift from Didier Trono, Addgene). After 3 d incubation, the culture medium was collected and concentrated five times by ultrafiltration (Amicon Ultra-50 with MWCO 100K, Millipore, Bedford, MA, USA). Viral titers were measured by a kit (Lenti-X p24 Rapid titer kit, #632200. Takara, Kusatsu, Japan). To suppress Mul1 expression, Mul1 shRNA lentiviruses (shMul1) were prepared using commercial plasmids (TRCN0000040742 and TRCN0000328514, Sigma); TRCN0000040742, 5′-CCGGCCTCTTCTTCATCCTGAGGAACTCGAGTTCCTCAGGATGAAGAAGAGGTT TTTG-3′ and TRCN0000328514, 5′-CCGGCTGTCGCTTCAGGAGCATAAGCTCGAGCTTATGCTCCTGAAGCGACAGTT TTTG-3′. TRCN0000040742 and TRCN0000328514 were used for the main experiments and validation of Mul1 knockdown, respectively. Infection of J774A.1 cells with lentiviruses was performed at 50% confluency in a 10 cm-dish with 8 μg polybrene (Sigma) in the absence of FBS. One day after viral infection, the culture medium was replaced with fresh RPMI1640 containing 10% FBS. Plasmids were transfected into RAW264.7 cells with FuGENE HD (Promega, Madison, WI, USA). At 50 % confluency, the culture medium was replaced with fresh RPMI1640 containing 10% FBS, 15 μg of pMul1-FLAG and 15 μg of pAsc-Myc mixed with FuGENE HD at a ratio of 1:3 in 500 μl Opti-MEM (Thermo Fisher) were transfected, and culture medium was refreshed the next day.

### *In vivo* and *in vitro* Asc ubiquitination

Both *in vivo* and *in vitro* Mul1-mediated Asc ubiquitination assays were performed, based on our previous report [62]. For detection of Asc ubiquitination *in vivo*, HEK293 cells at 70% confluence in a 10 cm-dish were transfected with pMul1-FLAG (2 μg), pUbi-HA (2 μg), and pAsc-Myc (2 μg). After overnight incubation, the cells were treated with 5 μM MG132 for 16 h and lysed by sonication in 500 μl of Brij lysis buffer (10 mM Tris pH 7.5, 150 mM NaCl, 2.5 mM EDTA, 0.875% Brij 97, 0.125% NP40) containing protease and phosphatase inhibitor cocktails. After centrifugation at 18,000 × *g* at 4 °C for 20 min, Asc was immunoprecipitated overnight with 20 μl of anti-Myc agarose beads at 4 °C. After washing the beads, bound Asc was eluted by boiling in 5× Laemmli sample buffer and ubiquitinated Asc in western blot was detected with anti-HA antibody. To detect Mul1-mediated Asc ubiquitination *in vitro*, the wild-type and mutant (C302/305R) recombinant Mul1 proteins were produced in *Sf9* cells using the Bac-to-Bac Baculovirus expression system (Invitrogen, Carlsbad, CA, USA) and purified by a nickel-chelating resin (Life Technologies, Carlsbad, CA, USA). *In vitro* ubiquitination of Asc was performed by incubating 300 ng of mouse recombinant Asc (CSB-EP861664MO, CUSABIO, Tx, USA), 1 µg of recombinant Mul1, 0.1 µg of mouse E1 (UBE1, Boston Biochem, Cambridge, MA, USA), and 1 µg of mouse E2 (UBE2E3, Boston Biochem) in 40 µl of reaction buffer (Boston Biochem) for 4 h at 30 °C. The reaction mixtures were then heated in 5× Laemmli sample buffer at 95 °C for 5 min, and the western blot was performed with anti-ubiquitin and anti-Asc antibodies.

### Co-immunoprecipitation

Binding between exogenous Mul1 and exogenous Asc was assessed by reciprocal co-immunoprecipitation analysis. HEK293 cells were transfected with 2 μg each of pMul1-FLAG and pAsc-Myc. Two days after transfection, the cells were treated with 5 μM MG132 for 16 h before harvesting the cells. After lysis in 500 μl of the Brij lysis buffer, cell lysates were centrifuged at 18,000 × *g* at 4 °C for 20 min. A total of 3.5 mg protein lysate was incubated with 60 μl of anti-Myc or anti-FLAG agarose beads overnight at 4 °C. After washing five times with Brij buffer, Mul1 bound to Asc or Asc bound to Mul1 was detected by western blotting with anti-Mul1 or anti-Asc antibody, respectively. For interaction of endogenous Mul1 and endogenous Asc, J774A.1 cells were treated with 100 ng/ml LPS and 5 μM MG132 for 6 h and stimulated with 5 mM ATP for 1 h. Thereafter, cell lysates were prepared, endogenous Asc from a total of 3.5 mg protein was immunoprecipitated with 1 μg of anti-Asc antibody and 60 μl of protein A/G agarose beads (Sigma), and bound Mul1 was detected with anti-Mul1 antibody by western blotting. For detection of endogenous Asc bound to Mul1, Mul1-FLAG was expressed in J774.A1 cells, because immunoprecipitation-compatible anti-Mul1 antibody was not commercially available. After transfection of pMul1-FLAG and treatment with LPS, ATP, and MG132 as above, Mul1-FLAG in a total of 3.5 mg protein was immunoprecipitated with 40 μl of anti-FLAG agarose beads, and bound Asc was detected with anti-Asc antibody by western blotting.

### Statistics

Data are expressed as means ± SD, and analyzed by standard one-way ANOVA or Kruskal-Wallis one-way ANOVA on ranks followed by Holm-Sidak or Student-Newman-Keuls methods for multiple comparisons after testing for normality using Shapiro-Wilks method using a software (SigmaPlot, Systat Software, San Jose, CA, USA). To estimate inhibition of Asc reduction by proteasome and autophagy inhibitors, data in the absence and presence of Mul1 overexpression were analyzed by one-tailed Student *t*-test after normality test by Shapiro-Wilks method.

## Acknowledgments

This research was supported by the National Research Foundation of Korea (NRF) funded by the Ministry of Education (NRF-2017R1D1A1B03028344, YS Lee; NRF-2017R1A6A3A11033581, J Kim).

## Author contributions

Study concept and design (J Kim, GY Seo, YS Lee); experiments and data analysis (J Kim, YS Lee); and preparation of manuscript (J Kim, Y Lee, GY Seo, YS Lee).

---

**Figure EV1.**
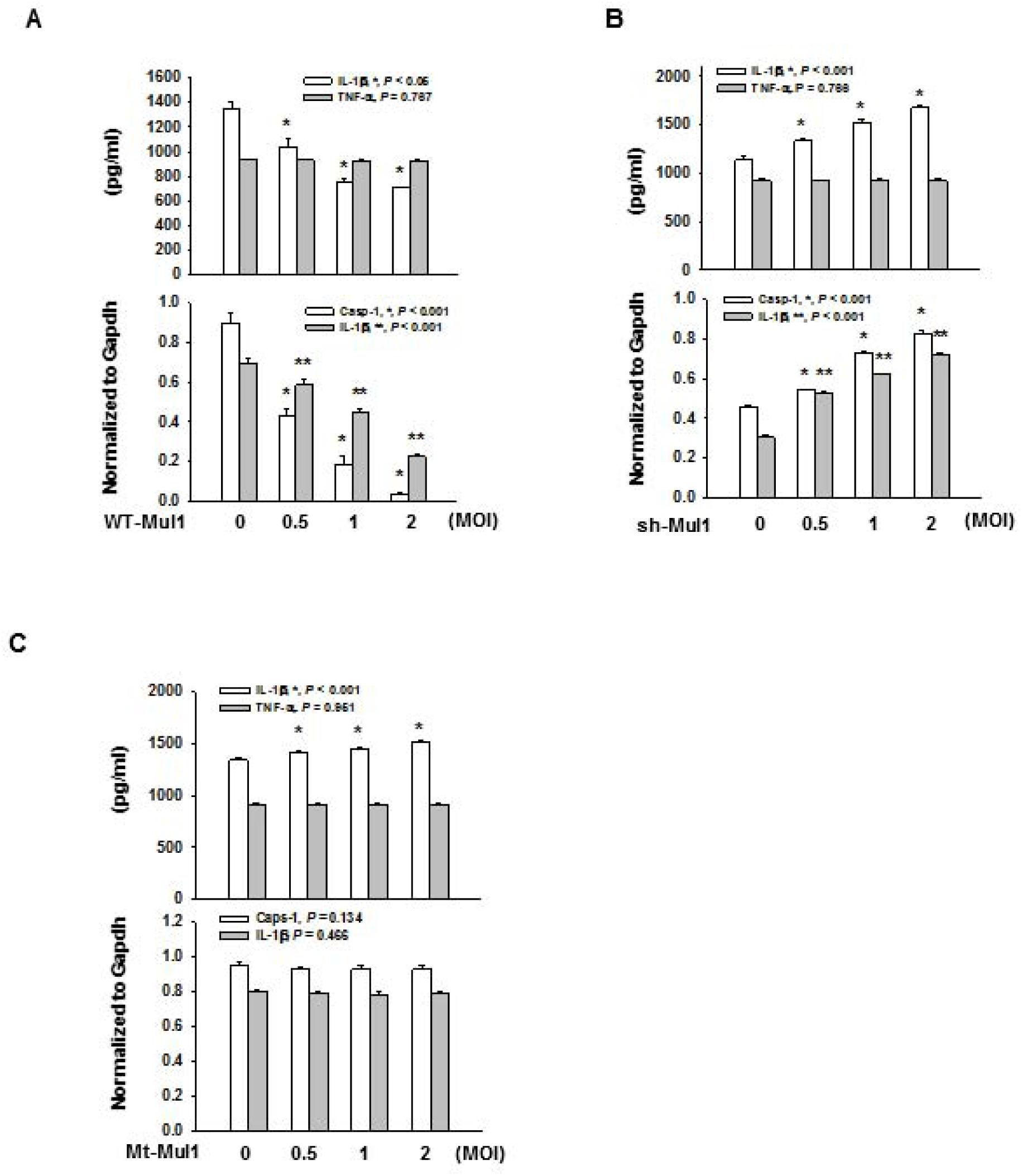
Mul1 negatively regulates the Nlrp3 inflammasome activation induced by LPS and nigericin. Experiments performed as in Figure 1, except Nlrp3 inflammasome activation induced by LPS and nigericin. For Nlrp3 inflammasome activation, LPS-primed J774A.1 cells were stimulated with 10 μM nigericin for 1 h. A Suppression of Nlrp3 inflammasome activation by Mul1 overexpression. * and ** indicates statistically significant difference, compared with the control lentivirus-infected group (n=4, * in IL-1β secretion by ANOVA on ranks and Student-Newman-Keuls method, * and ** in casp-1 (p20) and IL-1β (p17) by ANOVA and Holm-Sidak method). B Augmentation of Nlrp3 inflammasome activation by Mul1 knockdown. * and ** indicate statistically significant difference, compared with the control lentivirus-infected group (n=4, ANOVA and Holm-Sidak method). C Inhibitory effect of Mul1 on Nlrp3 inflammasome activation is dependent on its E3 ligase activity. * indicates statistically significant difference, compared with the control lentivirus-infected group (n=4, ANOVA and Holm-Sidak method)

**Figure EV2.**
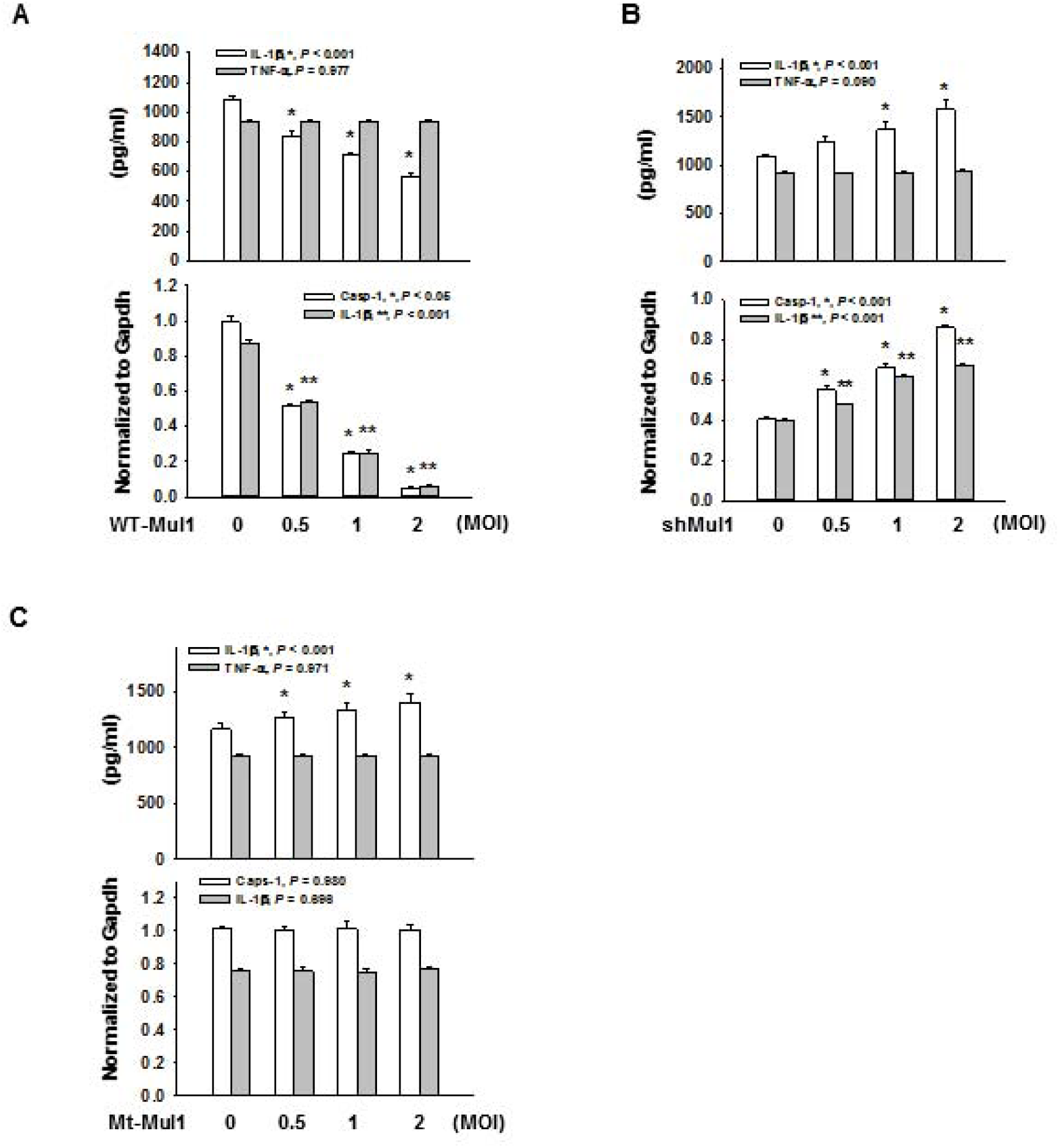
Mul1 negatively regulates the Nlrp3 inflammasome activation induced by LPS and alum. Experiments performed as in Figure 1, except Nlrp3 inflammasome activation induced by LPS and alum. For Nlrp3 inflammasome activation, LPS-primed J774A.1 cells were stimulated with 100 μg/ml alum for 1 h. A Suppression of the Nlrp3 inflammasome activation by Mul1 overexpression. * and ** indicate statistically significant difference, compared with the control lentivirus-infected group (n=4, * in IL-1β secretion and caspase-1 (p20) by ANOVA on ranks and Student-Newman-Keuls method, ** in IL-1β (p17) by ANOVA and Holm-Sidak method). B Augmentation of the Nlrp3 inflammasome activation by Mul1 knockdown. * and ** indicate statistically significant difference, compared with the control lentivirus-infected group (n=4, ANOVA and Holm-Sidak method). C Inhibitory effect of Mul1 on the Nlrp3 inflammasome activation is dependent on its E3 ligase activity. * indicates statistically significant difference, compared with the control lentivirus-infected group (n=4, ANOVA and Holm-Sidak method).

**Figure EV3.**
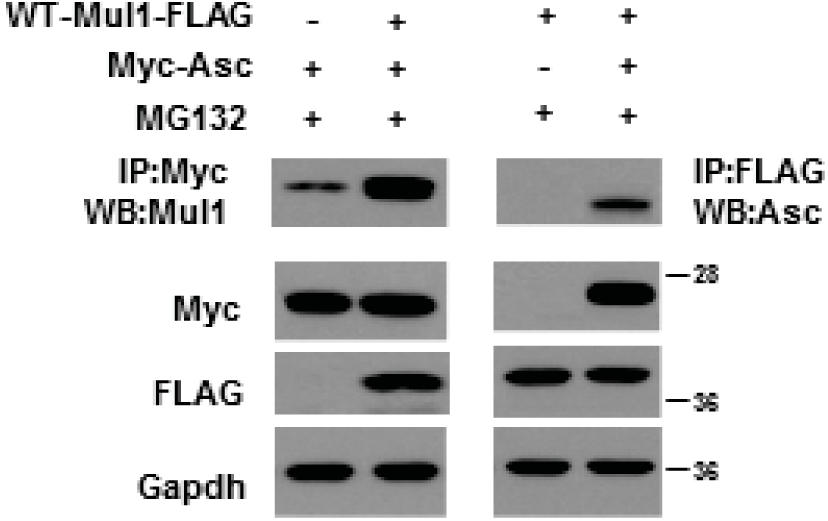
Interaction between exogenous Mul1 and exogenous Asc expressed in HEK293 cells. Mul1-FLAG and Asc-Myc were expressed in HEK293 cells and reciprocal co-immunoprecipitation assay was performed. *Left*; Mul1-FLAG was detected by co-immunoprecipitation with anti-Myc agarose beads, followed by western blotting with anti-Mul1 antibody. *Right*; Asc-Myc was detected by co-immunoprecipitation with anti-FLAG agarose beads, followed by western blotting with anti-Asc antibody.

